# Comparative study on the accuracy of speech recognition using a contact microphone attached to the surface of the head and neck

**DOI:** 10.1101/2024.04.17.589963

**Authors:** Takumi Asakura, Yuki Konuma

## Abstract

The accuracy of speech recognition through an air-conducted microphone can be less accurate under a highly noisy environment or when the volume of the user’s voice is relatively low. One solution to this problem is the use of contact microphones. However, neither the microphone locations that provide optimal speech recognition accuracy for each user nor the mechanisms underlying these contact forces have been clarified. In this study, we experimentally investigated the effects of placement, contact force, user gender, and speech recognition platform on the accuracy of speech recognition with contact microphones placed on the surface of the head and neck. The experimental results indicated that the mechanism underlying the influence of each factor on speech recognition accuracy differs for speech acquired at the neck and head locations. In particular, the effect of the user’s gender was significant for the neck-acquired sound, but not the head-acquired sound. The results also revealed that the microphone contact force did not affect the recognition accuracy or user discomfort for the head-acquired sound. Moreover, the results of speech recognition experiments in a simulated noisy environment showed that bone-conducted sounds acquired on the head and neck surfaces were more robust than air-conducted sounds.

## Introduction

It has been reported that as of 2023, at least 2.2 billion people globally are mildly or severely visually impaired [1]. Under such circumstances, many visually impaired people use electronic media such as smartphones to obtain information [2,3]. However, using the hands to input text may cause visually impaired people, especially those who use a cane or guide dog, to fall. In addition, it is difficult for such individuals to operate electronic media that relies on visual information. Therefore, there is a strong need to use applications without visual information. On the other hand, privacy protection in speech is important [4–8], and it is important to be able to perform speech recognition even under noisy environments using voices with as low a volume as possible. In addition, while there are many cases of healthy people using smartphones with voice control [9], there are also reports that many users are bothered by the voice of another uttered to control them [10]. Therefore, efficient speech recognition technology using low-volume voices is an important issue for not only physically challenged, but also healthy individuals. To acquire voice information efficiently, a contact microphone, which has the advantage of being less susceptible to ambient noise compared with an air-conducted microphone [11–13], can be used. Because bone-conducted speech acquired using a contact microphone and air-conducted speech acquired using an air-conducted microphone have different acoustic characteristics, some studies [13–15] have found that combining both types of microphones can improve the accuracy of speech recognition under noisy conditions. Previous studies have investigated the effects of the position of a contact microphone on the subjective intelligibility and quality of speech in the cases of bone-conducted sound obtained at the neck [16–18], at the head [19–21], and at both locations [22], and have confirmed that speech intelligibility and sound quality vary depending on the placement position of the contact microphone. On the other hand, generally available speech recognition systems are built based on air-conducted speech and thus not intended to be applied to the recognition of speech data acquired using contact microphones. Therefore, research has been conducted to improve the quality and accuracy of speech recognition for bone-conducted speech by signal processing [23–25]. In the field of clinical audiology, recommended contact forces have been specified [26,27] for the application of bone-conducting vibrators on the head. On the other hand, from the standpoint of perceiving bone-conducted sounds with a contact microphone, to our knowledge, no studies have investigated the optimal contact force. From the viewpoint of practical use, the accuracy of bone-conducted speech recognition by commonly available speech recognition systems has been examined [28], as have the effects of each factor, such as microphone placement on the neck, speech recognition model, and subject gender. However, that previous study was limited to the neck area only, and did not perform an overall verification of the speech recognition accuracy of bone-conducted sounds obtained using a contact microphone in the entire head and neck area.

Given the aforementioned background, the following research questions can be raised in the discussion of the more practical use of bone-conducted sounds obtained from the head and neck area:

1. What is the applicability of commonly available speech recognition systems based on the use of air-conducted microphones for speech recognition of bone-conducted sounds observed in the entire head and neck region?
2. How does the setting position of a contact microphone on the surface of the head and neck and the user’s physical characteristics influence the accuracy of speech recognition?
3. How does the contact force of the contact microphone against the skin affect the accuracy of speech recognition and the user’s discomfort?
4. How do bone-conducted sounds acquired on the surface of the head and neck under a noisy environment affect the accuracy of speech recognition compared with air-conducted sounds?
5. What comparative knowledge can be obtained regarding the pros and cons of the speech recognition performance of both head and neck bone-conducted sounds with respect to the various items mentioned above?

Given this background, the present study aimed to achieve efficient speech recognition in noisy environments using a contact microphone by comparing the speech recognition accuracy of bone-conducted sounds observed at each of the positions on the head and neck. The results of speech recognition for bone-conducted speech observed with contact microphones placed at seven locations on the head were compared with previous results for bone-conducted speech recognition acquired at eight locations on the surface of the neck [28]. Then, the effects of microphone position, subject gender, and microphone contact force were examined using three types of commonly available speech recognition software programs. In addition, the effects of three different contact forces on user discomfort were investigated in a subjective evaluation experiment. Finally, to obtain knowledge for a more practical bone-conducted sound recognition technique, recognition accuracy under a simulated acoustic environment with environmental noise in cities was discussed.

### Overview and flow of study

In this study, we evaluated the effects of differences in the placement of contact microphones placed on the surface of the head and neck on the accuracy of speech recognition. The data obtained from the surface of the head were compared with previous results of those from the neck [28]. For reference, we also acquired air-conducted speech data, which were obtained using a microphone for air-conducted sound measurement. These data were then compared with bone-conducted speech data to evaluate the effect of each speech acquisition method on recognition accuracy.

First, we conducted an experiment to investigate the accuracy of both bone- and air-conducted speech recognition under ideal conditions as Experiment I. Then, the main effects and interactions of microphone position, subject gender, and speech recognition system were discussed. When placing a contact microphone on the head, there is concern that the crimping of the microphone on the surface of the head may cause discomfort. Therefore, a subjective evaluation experiment of the annoyance caused by mounting the microphone on the head was conducted as Experiment II. Then, as Experiment III, to verify the recognition accuracy in a realistic situation, we simulated a noisy environment in a laboratory using background sounds recorded in an actual city environment and evaluated changes in speech recognition accuracy for both the contact and airborne microphones when the sound pressure level (SPL) of the background sound changes.

## Methods

In this section, the common settings of bone-conducted speech measurement and statistical analysis through experiments I, II, and III are described, along with the details of each experiment.

### Evaluation of bone-conducted speech recognition

The following three commonly available systems were used for speech recognition: AmiVoice (Advanced Media, Tokyo, Japan), Speech-to-text (Google, Mountain View, CA, USA), and Speech-to-text (Microsoft Azure, Microsoft, Redmond, WA, USA). These systems are henceforth referred to as Systems I, II, and III, respectively. We input speech information to these software systems through microphones, calculated the character error rate (CER) of the output sentences, and used this parameter to evaluate the recognition accuracy. These systems output recognized sentences by inputting sound files recorded in the manner described in the following section. As described in the following section, all speech data were recorded monaurally with a sampling frequency of 44.1 kHz and a quantization bit rate of 16 bit, and input to all speech recognition software in this format. Therefore, there was no difference in the resolution of the audio between the different software systems.

Next, the contact microphones were placed at eight positions on the throat [28], as shown in Fig 1(a), and at seven locations on the head, as shown in Fig 1(b). The installation positions on the throat were as described in a previous study [28]: upper thyroid cartilage (P1, P3, P5, P7), two vertical positions on the cricothyroid ligament (P2, P4, P6, P8), frontal positions (P1, P2), adjacent positions to the protruding cartilage (P3, P4), positions between P3 and P7 and between P4 and P8 (P5, P6), and anteroposterior positions of the mandibular angle (P7, P8). On the other hand, the installation positions on the head were the top of the head, forehead, temple, mandibular condyle, mandibular angle, mastoid, and chin. To simulate the use of the microphone in a real environment, the mounting positions were presented to the subjects orally and with the use of diagrams as appropriate, and the subjects were asked to put the device on themselves. Note that in this experiment, measurements were taken only on the right side of the body.

**Fig. 1.**
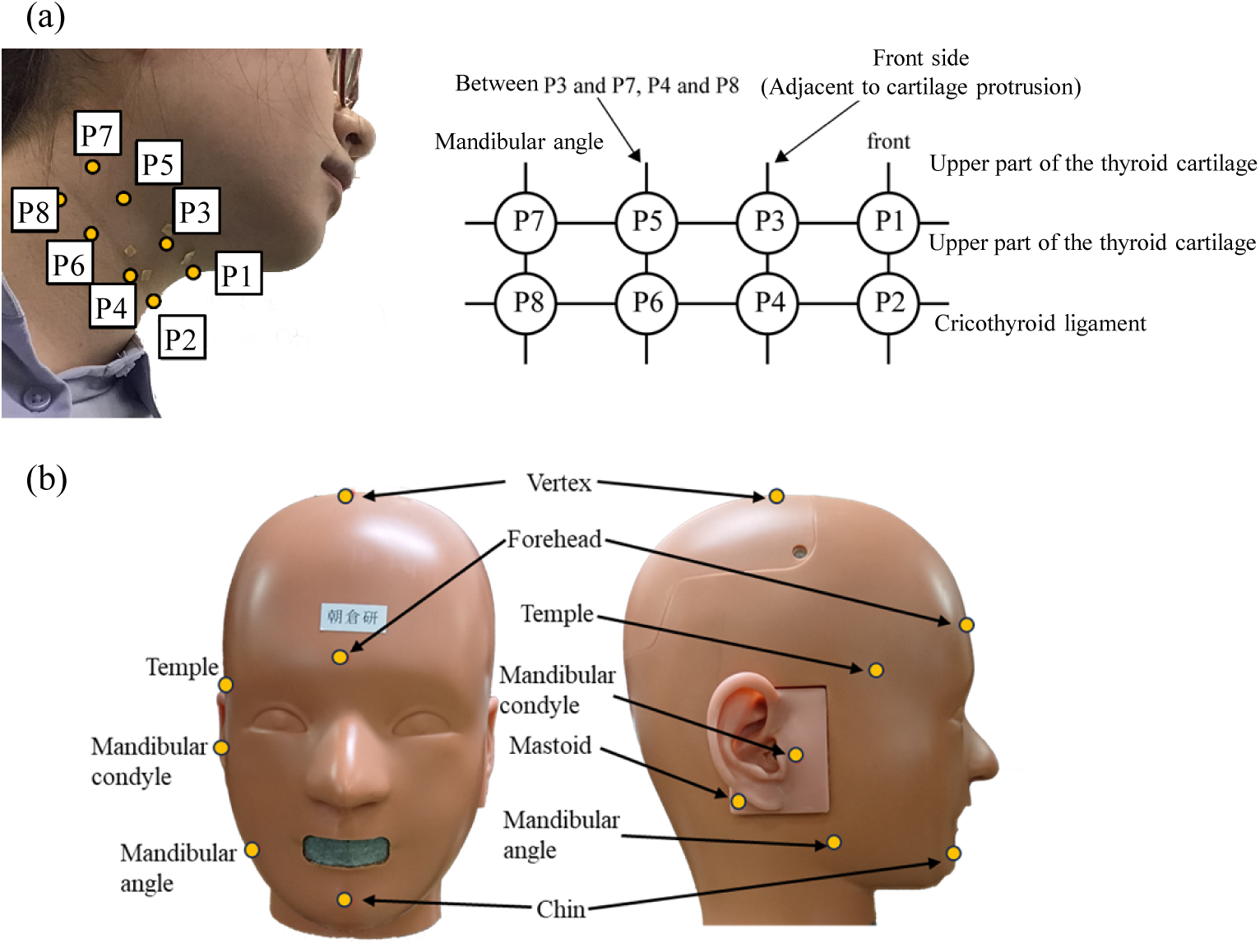
Mounting position of the contact microphone on the surface of the (a) neck and (b) head, respectively.

### Statistical analysis

Multi-way analysis of variance (ANOVA) was conducted to investigate the effects of each factor in relation to the gender, speech recognition software, microphone position, and contact force, and Dunnett’s multiple comparison test was used to assess the difference between each of the levels in each factor with significant main effects. Because normality could not be ensured, Friedman’s test was conducted in the multi-way ANOVA of the survey results on subjective impressions of discomfort. All statistical analyses were conducted using JMP (JMP Statistical Discovery LLC, Cary, NC, USA), with p values < 0.05 considered statistically significant.

### Basic characteristics of bone-conducted speech recognition observed in the neck and head (Experiment I)

The recording scheme for the speech data on the neck is described as follows. The study participants were five males and five females in their 20s (mean age: 22.8 years, standard deviation [SD]: 0.6 years). The equipment used for the recordings consisted of a condenser microphone (ECM-SP10; Sony, Tokyo, Japan) for air-conducted speech, a contact microphone (C9038A; RETEVIS, Ysair Technology, Shenzhen, China) for bone-conducted speech, and a smartphone (XperiaZ2 SO-03F; Sony). All recordings were monaural at a sampling frequency of 44.1 kHz with a quantization bit rate of 16 bit. The subjects were asked to speak three sentences, five times each, for a total of 15 sentences. The sentences were taken from parallel100 of the JVS corpus [29]. The equivalent A-weighted SPL during speech was set to 61.5–63.5 dB to avoid large variations of reproduced sound from subject to subject. The speech data were collected twice with at least a 1-week interval between each recording.

Next, the recording of speech data on the head was conducted using the same 10 subjects as the those in the neck recordings. The recording equipment used included a condenser and a contact microphone, same as those used for the neck measurement. Note that the data acquisition was conducted using an audio interface (Steinberg UR444C; Yamaha, Hamamatsu, Japan) instead of the smartphone used in the recording for the neck. The number of quantization bits and the sampling rate were the same in both cases, and there was no difference in the quality of the audio. The contact microphone was fixed to the neck using a neckband, as shown in Fig 2(a), but the contact microphone was fixed to the head using a three-dimension-printed fixation part and a bandage (KB25F; Nichiban, Tokyo, Japan), as shown in Fig 2(b). The contact force of the microphone was set to three conditions—2 N, 3 N, and 4 N—in reference to a previous discussion on bone-conducting vibrators [26]. A digital force gauge (FGJN-5; NIDEC, Kyoto, Japan) was used to measure the contact force. The string attached to the fixing part of the microphone was put on the hook of the force gauge, which was then pulled away slowly, and the measurement value was read when the microphone was lifted slightly from the skin. The contact force during the experiment was adjusted by tightening the bandage until it reached ±0.5 N of the target value in each condition. Note that the neckband originally attached to the microphone was cut off before use in the case of the head measurement. The audio settings in the recording were the same as those for the throat measurement for the part.

**Fig. 2.**
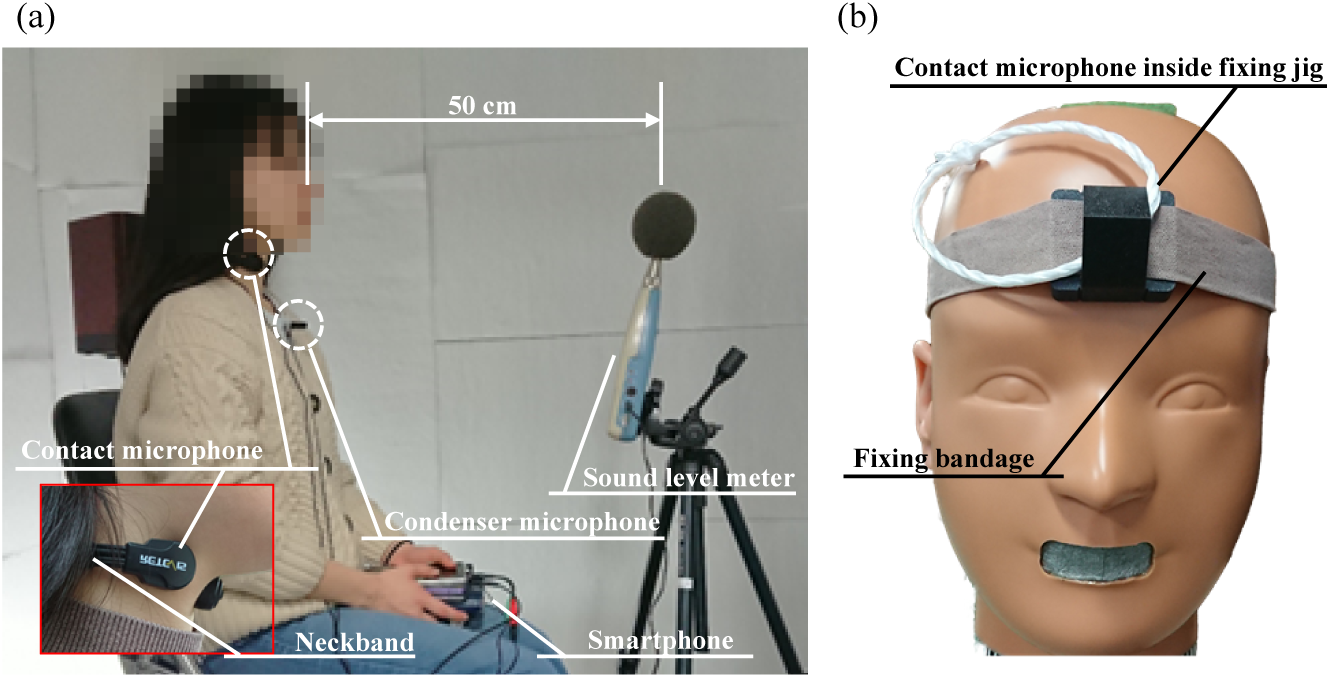
(a) Spatial relationship between the contact and condenser microphones and the sound level meter for monitoring the sound pressure level of the subject’s voice. (b) Setting condition of the contact microphone with a fixing jig and bandage on the surface of the head.

### Subjective annoyance with wearing a contact microphone (Experiment II)

Any pain, suffering, or discomfort that may have been caused by wearing the contact microphone was investigated in a subjective evaluation experiment. The subjects were 20 males and 10 females in their 20s (mean age: 23.3 years, SD: 0.9 years), and included the same five males and five females from the recognition accuracy experiment. The subjects were asked to rate their responses on a seven-point Likert-type scale ranging from “strongly agree” (score: 7) to “strongly disagree” (score: 1) (Fig 3).

**Fig. 3.**
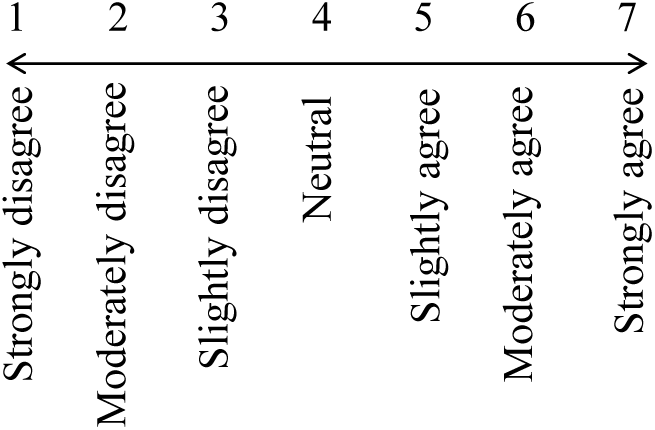
The seven-point Likert-type scale used in the subjective evaluation experiment.

### Bone-conducted speech recognition in a noisy environment (Experiment III)

In this experiment, both speech and background sound data were recorded separately and superimposed to create speech data that simulated speech in a realistic environment with background noise. The synthesized speech data with background sounds were then input to each speech recognition system, and the CER was calculated. The background sounds were recorded using a binaural microphone (BME-200; Adphox, Tokyo, Japan) on a street in Kashiwa city, Chiba, Japan. Three sets of 120 seconds of data were recorded, with equivalent A-weighted SPLs of 66.5, 66.5, and 69.4 dB, respectively. As these levels did not change much, we assumed that the background sounds at this area in the city were in a quasi-steady state. The steady parts were extracted from the waveforms, which were not mixed with specific sounds such as human voices and automobile sounds, and the extracted sounds were reproduced through four loudspeakers (NS-B700; Yamaha, Tokyo, Japan) with equivalent A-weighted SPLs at the test subject’s position of 65 dB and 70 dB (Fig 4). Then, background sounds were recorded using the air-conducted and contact microphones mounted in the same way as previously described. The positions used in this experiment were P5 and P7 on the throat, and the forehead and chin on the head. Then, speech data recorded in a real environment were simulated by superimposing the air- and bone-conducted sound data recorded when speech was uttered using the method described above and the background sound data recorded as air- and bone-conducted sound. The synthesized speech data were finally input to a speech recognition system, and the CER was calculated from the recognition results.

**Fig. 4.**
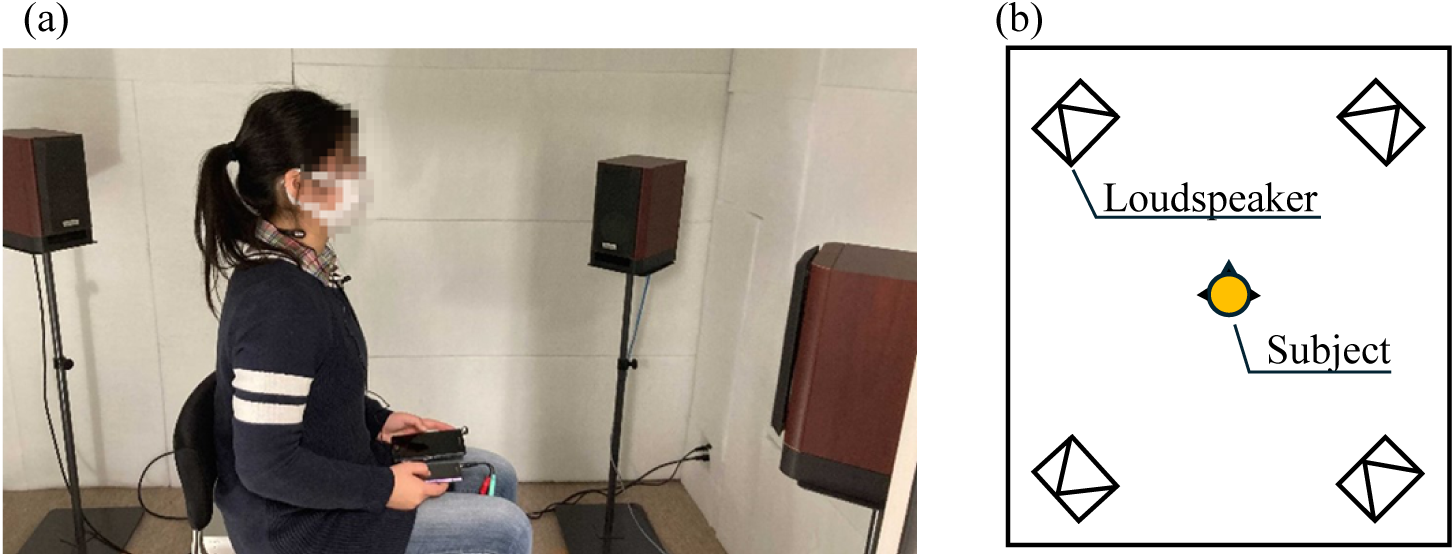
(a) Recording situation of air- and bone-conducted background sounds and (b) the spatial relationship between the four loudspeakers and the evaluation point inside the room.

## Results

### Experiment I

Fig 5(a) to (b) [28] and (c) to (h) show the CERs obtained for the throat and head, respectively. Specifically, Fig 5(a) and (b) summarizes the CERs of the air- and bone-conducted speech data obtained by each of the speech recognition systems for each mounting position, while Fig 5(c) to (h) summarizes the CERs of the air- and bone-conducted speech data obtained under each contact force and speech recognition system condition for each mounting position.

The results indicated that the air-conducted speech had a low CER in all cases. For the bone-conducted speech, the results obtained from the head showed a lower CER than those from the throat. Fig 5(b) shows that the bone-conducted speech obtained from the throat had a lower CER at positions P5–P8 than at positions P1–P4 for all speech recognition systems. In addition, the CER was generally lower at the odd compared with the even number positions, i.e., at the height of the upper thyroid cartilage compared with the cricothyroid ligament.

Regarding the results obtained from the head, Fig 5(f) shows that the CER in System III was lower at all forces and positions. By contrast, the CERs in Systems I and II differed by position: the CERs in the forehead and chin were relatively low in System I, whereas those in the temples were higher than in the other positions in System II.

**Fig. 5.**
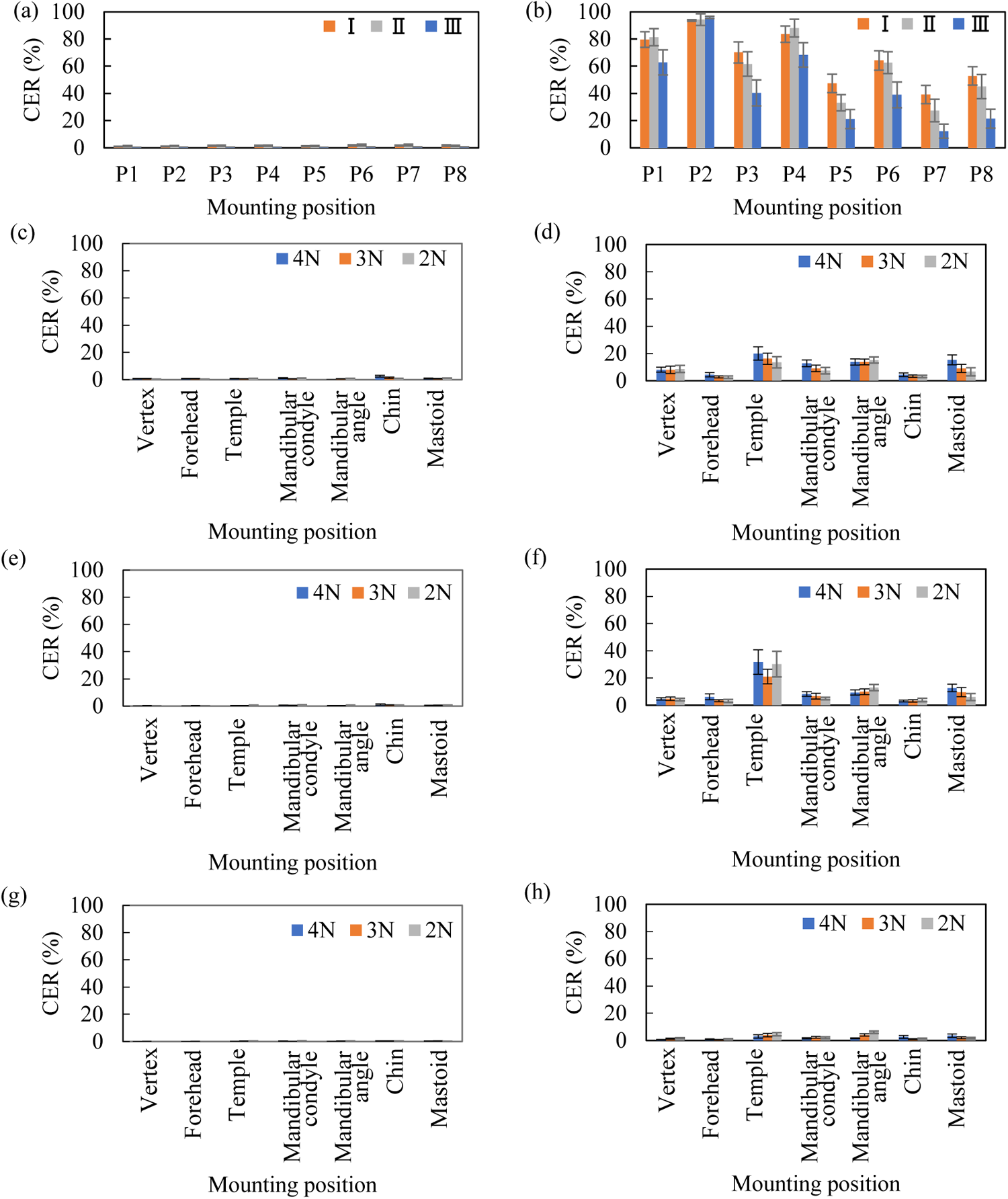
(a, b) Results of character error rates (CERs) [28] evaluated by Systems I, II, and III from the signals obtained by (a) the condenser microphone and (b) the contact microphone on the surface of neck. (c–h) Results of CERs evaluated from the signals obtained by (c, e, g) the condenser microphone and (d, f, h) the contact microphone using (c, d) System I, (e, f) System II, and (g, h) System III, respectively.

### Experiment II

Fig 6 shows the subjective results of the pain, suffering, and discomfort experienced when wearing the microphone on the head. As shown in the figure, the subjective scores for pain, suffering, and discomfort tended to decrease with increasing contact force, but this trend varied in each of the mounting positions. The subjective scores for the temples and forehead tended to be relatively high for all contact forces.

**Fig. 6.**
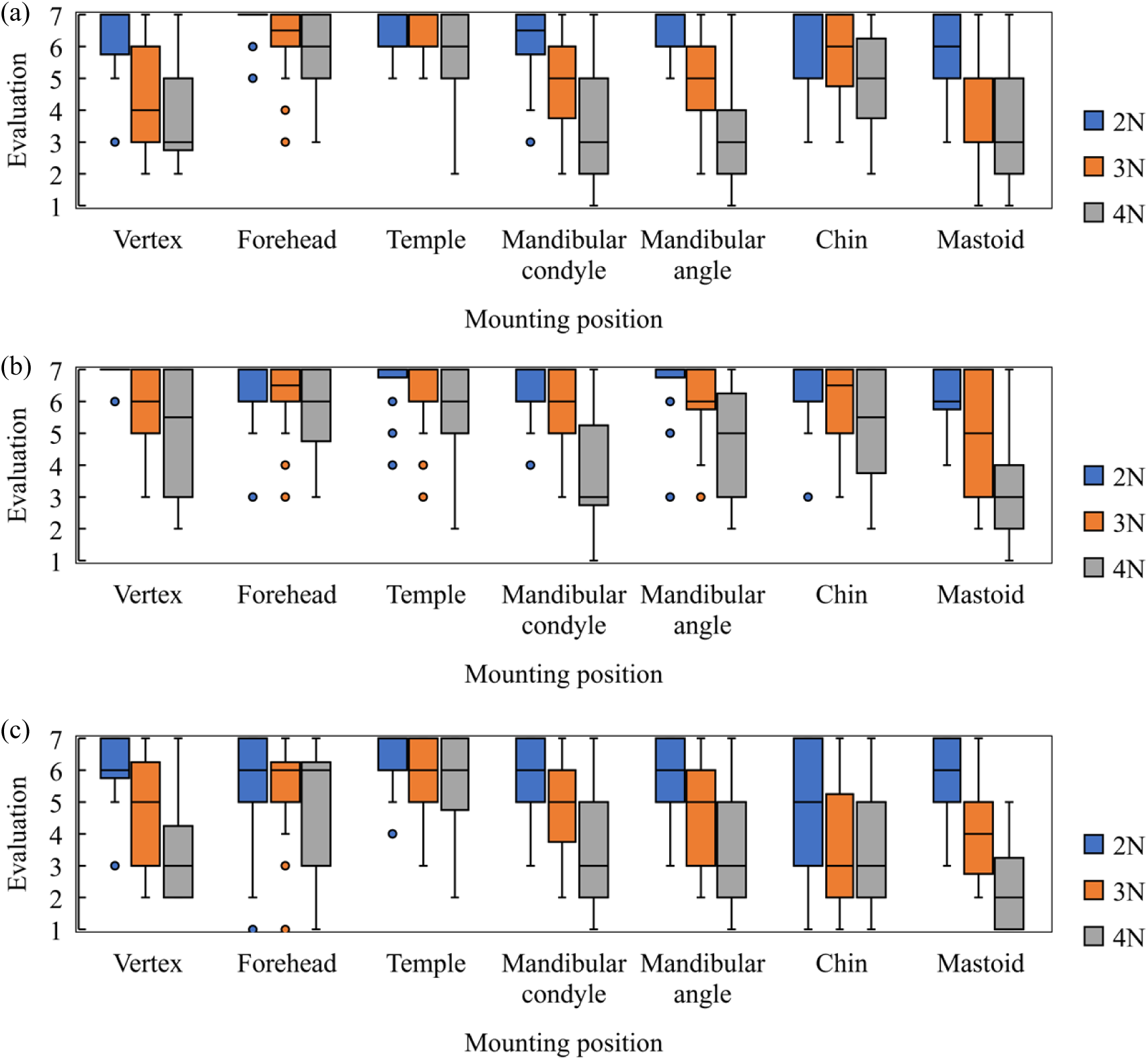
Results of the subjective scores for (a) pain, (b) suffering, and (c) discomfort when the contact microphone was mounted at each position on the head.

### Experiment III

The CERs obtained at P5 and P7 on the throat are shown in Fig 7(a), and those obtained at the forehead and chin are shown in Fig 7(b–d). As shown in Fig 7(a), the CER of air-conducted speech was the smallest under the background sound of 65 dB. However, under the background sound of 70 dB, the CERs of both air- and bone-conducted speech were similar, and in some conditions, the CER of bone-conducted speech was lower than that of air-conducted speech. Fig 7(b–d) shows that the CERs of bone-conducted speech were the same or lower than those of air-conducted speech under all conditions for the forehead and chin, the CERs increased with increasing background SPL only for the forehead.

**Fig. 7.**
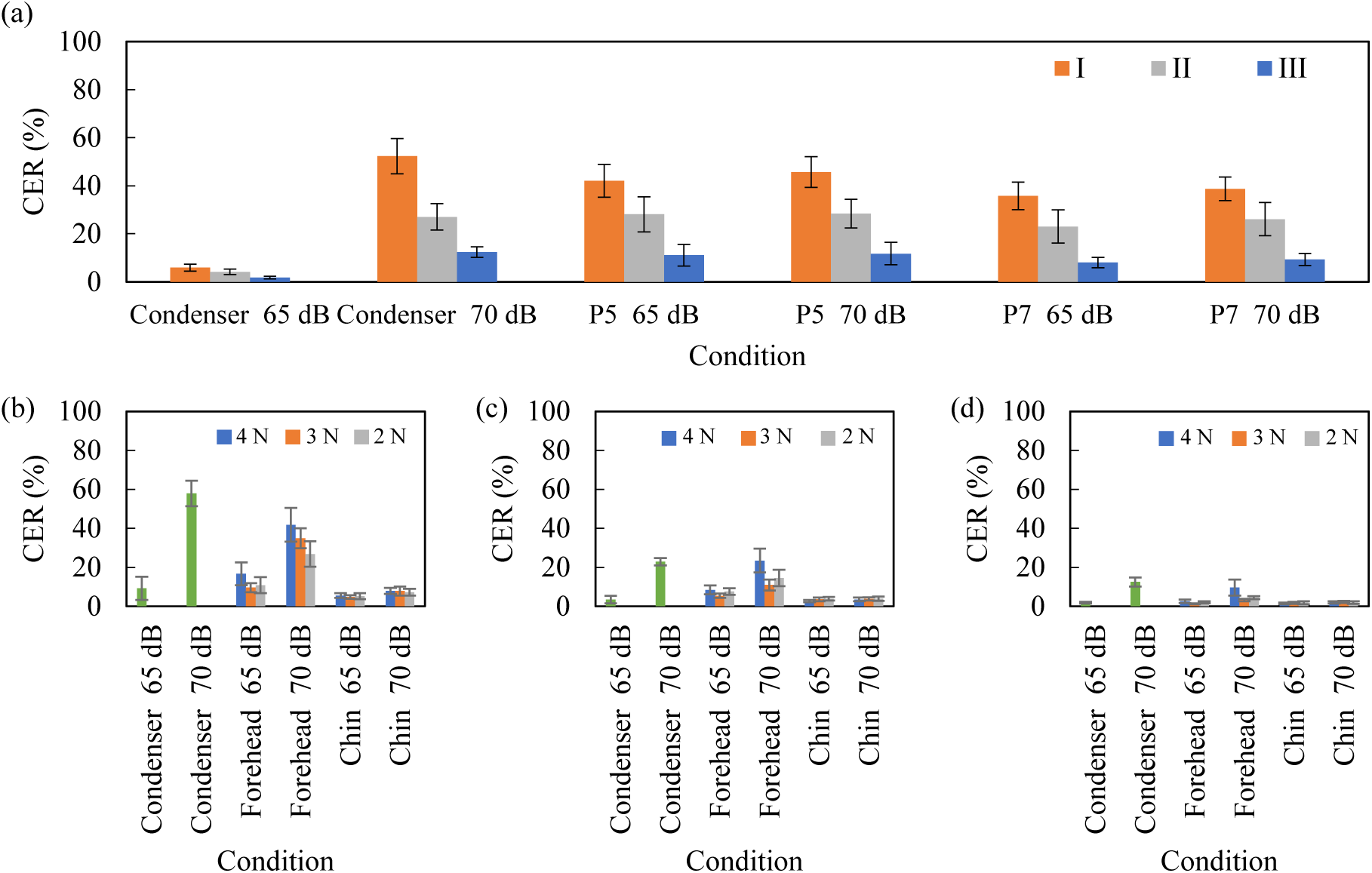
(a) The character error rates (CERs) obtained from the P5 and P7 signals on the neck as evaluated by Systems I, II, and III. (b–d) The CERs obtained from the forehead and chin signals as evaluated by (b) System I, (c) System II, and (d) System III.

## Discussion

### Effect of each factor on bone-conducted speech recognition

Table 1 shows the results [28] of the ANOVA conducted for the CERs obtained from the contact microphones as three factors: speech recognition system, mounting position, and gender. No significant differences were found for the second-order interactions, but significant differences were found for the main effects and first-order interactions for all three factors.

**Table 1.** Results of an analysis of variance [28] conducted for the CERs obtained from the bone-conducted signals on the neck (\**p* < 0.05, \*\**p* < 0.001).

| Factor | Sum of squares | df | Mean square | F-value | P-value | Significance |
| --- | --- | --- | --- | --- | --- | --- |
| Gender | 31,411 | 1 | 31,411 | 6.6 | <0.05 | * |
| Mounting position | 274,072 | 7 | 39,153 | 98.6 | <0.001 | ** |
| ASR system | 50,605 | 2 | 25,303 | 63.7 | <0.001 | ** |
| ASR system - Gender | 49,24 | 2 | 2,462 | 6.2 | <0.05 | * |
| Mounting position - Gender | 26,858 | 7 | 3,837 | 9.7 | <0.001 | ** |
| ASR system - Mounting position | 13,616 | 14 | 973 | 2.4 | <0.05 | * |
| ASR system - Gender - Mounting position | 4,774 | 14 | 341 | 0.9 | 0.604 | NS |

On the other hand, Table 2 shows the results of a four-way ANOVA on the CER of bone-conducted speech obtained from the heady using the following four factors: speech recognition system, mounting position, gender, and contact force. The results showed that the main effects of the speech recognition system and mounting position and their interactions were significant, whereas the main effects of gender and contact force were not. The interaction between the speech recognition system and mounting position is shown in Fig 8. In this figure, the installation positions on the horizontal axis are sorted in descending order of the CER.

**Table 2.**
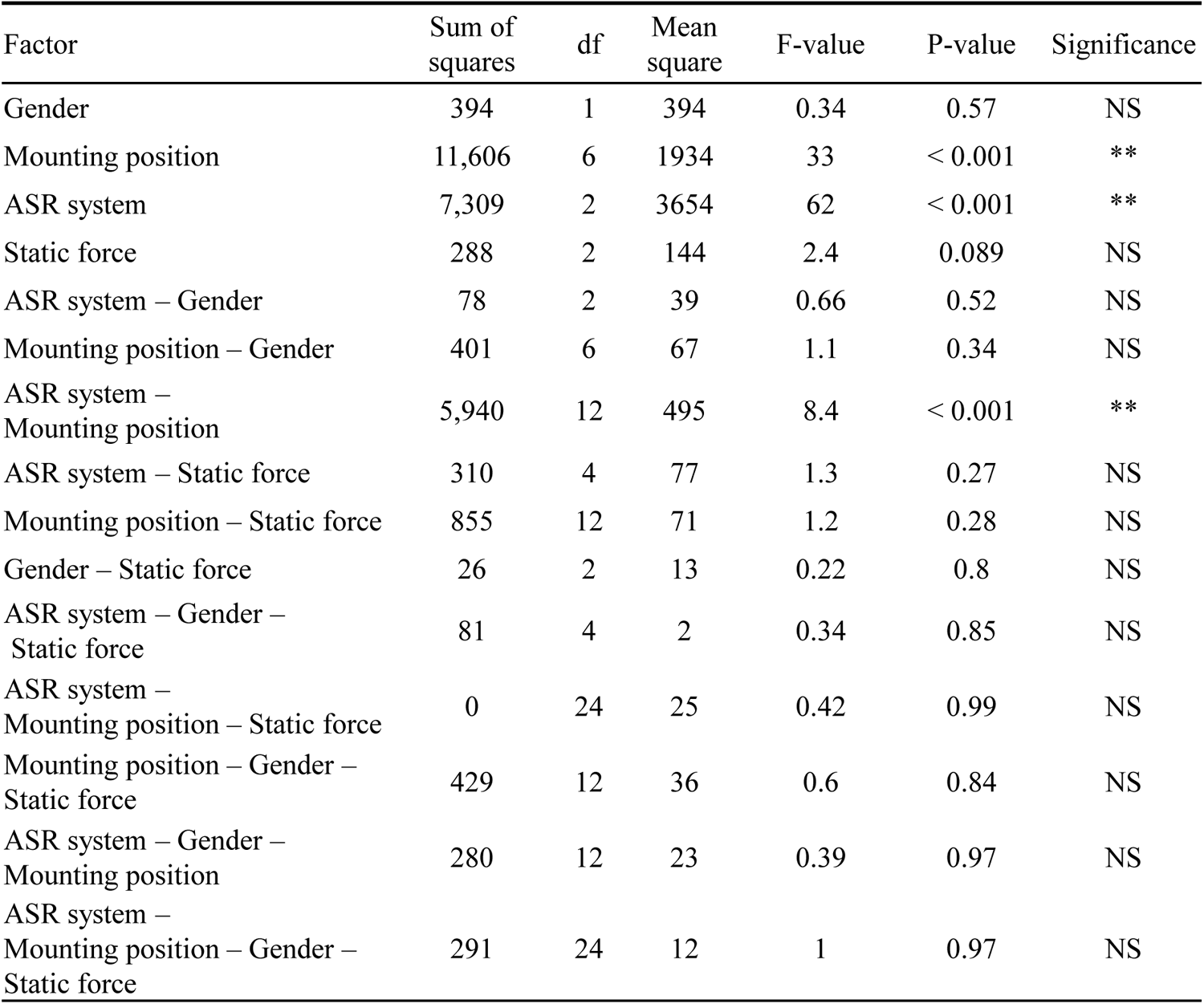
Results of an ANOVA for the CERs obtained from the bone-conducted signals on the head (\**p* < 0.05, \*\**p* < 0.001).

| Factor | Sum of squares | df | Mean square | F-value | P-value | Significance |
| --- | --- | --- | --- | --- | --- | --- |
| Gender | 394 | 1 | 394 | 0.34 | 0.57 | NS |
| Mounting position | 11,606 | 6 | 1934 | 33 | < 0.001 | ** |
| ASR system | 7,309 | 2 | 3654 | 62 | < 0.001 | ** |
| Static force | 288 | 2 | 144 | 2.4 | 0.089 | NS |
| ASR system – Gender | 78 | 2 | 39 | 0.66 | 0.52 | NS |
| Mounting position – Gender | 401 | 6 | 67 | 1.1 | 0.34 | NS |
| ASR system – Mounting position | 5,940 | 12 | 495 | 8.4 | < 0.001 | ** |
| ASR system – Static force | 310 | 4 | 77 | 1.3 | 0.27 | NS |
| Mounting position – Static force | 855 | 12 | 71 | 1.2 | 0.28 | NS |
| Gender – Static force | 26 | 2 | 13 | 0.22 | 0.8 | NS |
| ASR system – Gender – Static force | 81 | 4 | 2 | 0.34 | 0.85 | NS |
| ASR system – Mounting position – Static force | 0 | 24 | 25 | 0.42 | 0.99 | NS |
| Mounting position – Gender – Static force | 429 | 12 | 36 | 0.6 | 0.84 | NS |
| ASR system – Gender – Mounting position | 280 | 12 | 23 | 0.39 | 0.97 | NS |
| ASR system – Mounting position – Gender – Static force | 291 | 24 | 12 | 1 | 0.97 | NS |

**Fig. 8.**
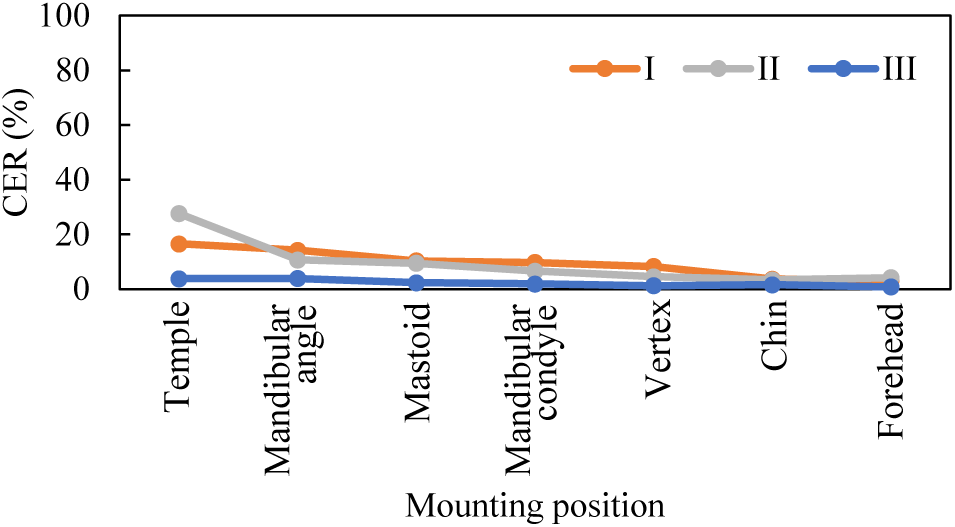
The character error rates (CERs) obtained at each mounting position on the head under each of the conditions of the speech recognition systems.

A simple main effect was obtained by conducting a post hoc test, and the results showed an effect of mounting position for Systems I and II, but no effect for System III (System I: *p* < 0.001, System II: *p* < 0.001, System III: *p* = 0.619). No effect of the speech recognition system on the CER was found for the forehead and chin (forehead: *p* = 0.544, chin: *p* = 0.211). Because an effect of the mounting position was observed, a multiple comparison of the CERs obtained by Systems I and II between each of the mounting positions was conducted, with the results shown in Fig 9. No significant difference was found between the forehead and chin for any of the speech recognition systems. Therefore, lower CERs were obtained for the forehead and chin regardless of the other factors of speech recognition system and gender.

A comparison of the effects of various factors on the speech recognition of bone-conducted sounds acquired in the neck and head revealed that the effect of gender was only observed in the neck; this effect was not observed in the bone-conducted sounds acquired from various parts of the head. A previous study that examined whether bone-conducted sounds could be detected when a bone-conducted vibrator was worn on the head, neck, and back found that males with a lower body fat percentage could detect sounds at a more distant mounting position, but no relationship was found between body fat percentage and auditory perception in females [30]. In other words, gender differences have been identified, as has a relationship between body fat percentage and hearing. It is possible that the gender differences observed in this study are also attributable to differences in body fat percentage between men and women. However, although there is a tendency for sound transmission to occur more efficiently with a lower body fat percentage, as in the aforementioned paper, there are cases in which females are noted to have more subcutaneous fat than males [31] and, conversely, males are noted to have more fat in the neck area [32].

These previous findings make a clear causal relationship in the present study difficult to elucidate. Furthermore, the neck is close to the vocal cords and has complex musculature. In addition, there is a possibility that differences in skeletal shape due to gender, such as differences in the degree of development of the throat, might have a combined effect on recognition accuracy. On the other hand, because the head surface has less subcutaneous fat and more contact with bone than the neck, it is natural that no gender differences were seen on the head.

**Fig. 9.**
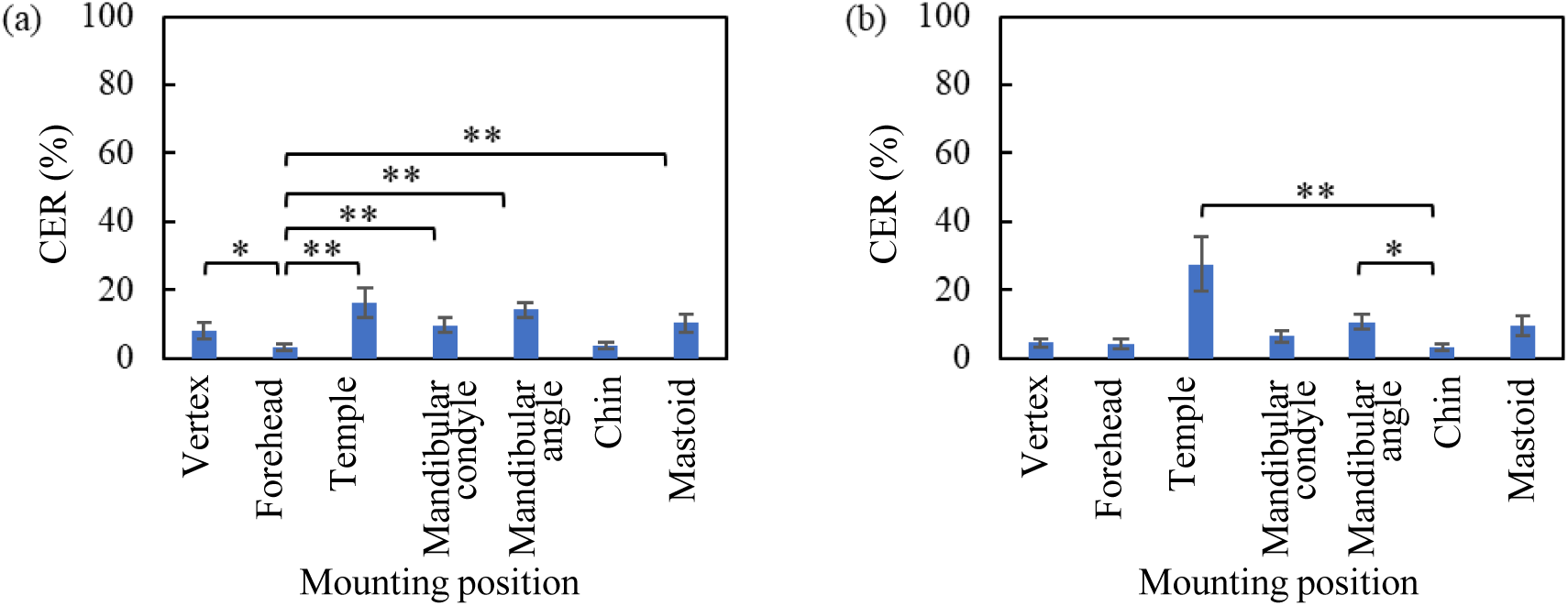
Results of multiple comparisons of character error rates (CERs) between each of the mounting position conditions in (a) System I and (b) System II, respectively.

### Subjective annoyance from wearing the bone-conducted microphone

Table 3 shows the results of a Friedman’s two-way ANOVA on the subjective scores regarding pain, suffering, and discomfort, with the two factors of contact force and mounting position. As shown in the table, no significant differences in pain, suffering, or discomfort by contact force and mounting position were observed. However, considering the fact that many respondents stated that it was “difficult to speak” and the chin was “in the way when speaking” in the free description column, the forehead may be more suitable for practical use in bone-conducted speech recognition compared with the chin.

**Table 3.** Results of Friedman’s test for the subjective scores for (a) pain, (b) suffering, and (c) discomfort.

| (a) |  |  |  | (b) |  |  |  | (c) |  |  |  |
| --- | --- | --- | --- | --- | --- | --- | --- | --- | --- | --- | --- |
| Factor | $\chi^2$ | P-value | Significance | Factor | $\chi^2$ | P-value | Significance | Factor | $\chi^2$ | P-value | Significance |
| Mounting position | 7.57 | 0.27 | NS | Mounting position | 12.17 | 0.06 | NS | Mounting position | 10.30 | 0.11 | NS |
| Static force | 2.42 | 0.30 | NS | Static force | 2.57 | 0.28 | NS | Static force | 2.16 | 0.34 | NS |

Tran et al. [26] and McBride et al. [27, 28] subjectively evaluated the intelligibility and quality of bone-conducted speech recorded at positions on the head, and both studies found that the forehead was best in terms of intelligibility and quality. They also stated that this position should be avoided because the subjective evaluation of intelligibility and quality obtained near the chin was relatively low owing to the lack of stable contact between the microphone and skin during speech, which differs from the low CER observed at the chin in the present experiment. The subjective evaluation of speech quality in those previous studies was lower because the subjects heard a noise caused by the friction between the contact microphone and the skin during speech, whereas in the present study, the friction sound was filtered out by noise reduction, so it did not affect the recognition accuracy.

### Bone-conducted speech recognition in a noisy environment

To discuss the effects of background sounds on speech waveforms, signal-to-noise (SN) ratios for each condition are shown in Fig 10. The SN ratio was calculated as the ratio of SPLs of the speech to the background sounds. The SN ratio of the bone-conducted speech was higher than that of the air-conducted speech at all positions. The SN ratios at P5, P7, and the chin were similar, while the forehead had a lower SN ratio than these locations. Besides, in the previous results shown in Fig 7, the CERs at P5, P7 and the chin did not change as a result of the increased SPL of background sounds, while those at the forehead were largely reduced. An explanation for this is that the SN ratio decreased as the background SPL increased, but the extent of the decrease was not sufficiently large to affect speech recognition. On the other hand, the SN ratio of the forehead was considered to be sufficient to cause a decrease in recognition accuracy.

The relative SPLs of the bone-conducted speech and background sound signals at each mounting position is shown in Fig 10. As seen in Fig 10(a), the SPLs of the background sounds on the forehead and chin were similar. However, Fig 10(b) shows that the forehead had a smaller speech signal than P5, P7, and the chin. From this result, it was estimated that the SPL of the bone-conducted speech signal detected at the forehead was small, and therefore, unlike other positions, the CER increased as a result of the increase of the SPL of the background sound.

For the above reasons, the CER of the bone-conducted speech obtained at the chin was lower than that of the air-conducted ones under all conditions. In addition, the change in the CER with the increased SPL of the background sounds was small, so in commonly available speech recognition systems, the CERs of bone-conducted speech signals are robust to background noise compared with air-conducted speech signals. However, it should be noted again that, as mentioned in the previous section, the attachment of the microphone to the chin received negative comments from the viewpoint of ease of speaking. Therefore, the contact microphone is useful in the daily life environment where the background noise level is high.

Because the change in the CER with respect to the change in the SPLs of background sounds was small in P5 and P7 on the throat, the contact microphone can be said to be generally useful in daily life, where background sounds are relatively intense. On the other hand, the forehead could be a useful microphone mounting position if it is possible to apply a microphone casing that can reduce the effects of background sounds.

**Fig. 10.**
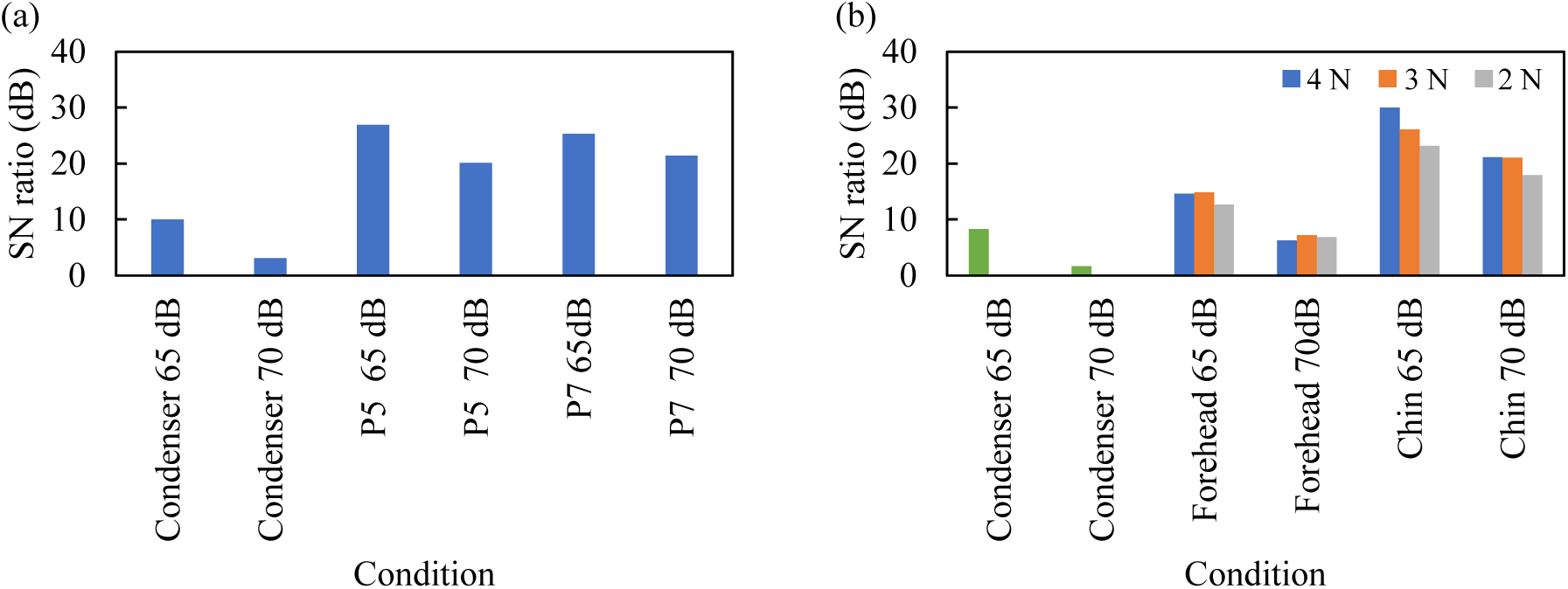
Signal-to-noise (SN) ratios for each of the mounting position and reproduced sound pressure level conditions for the (a) neck and (b) head, respectively.

**Fig. 11.**
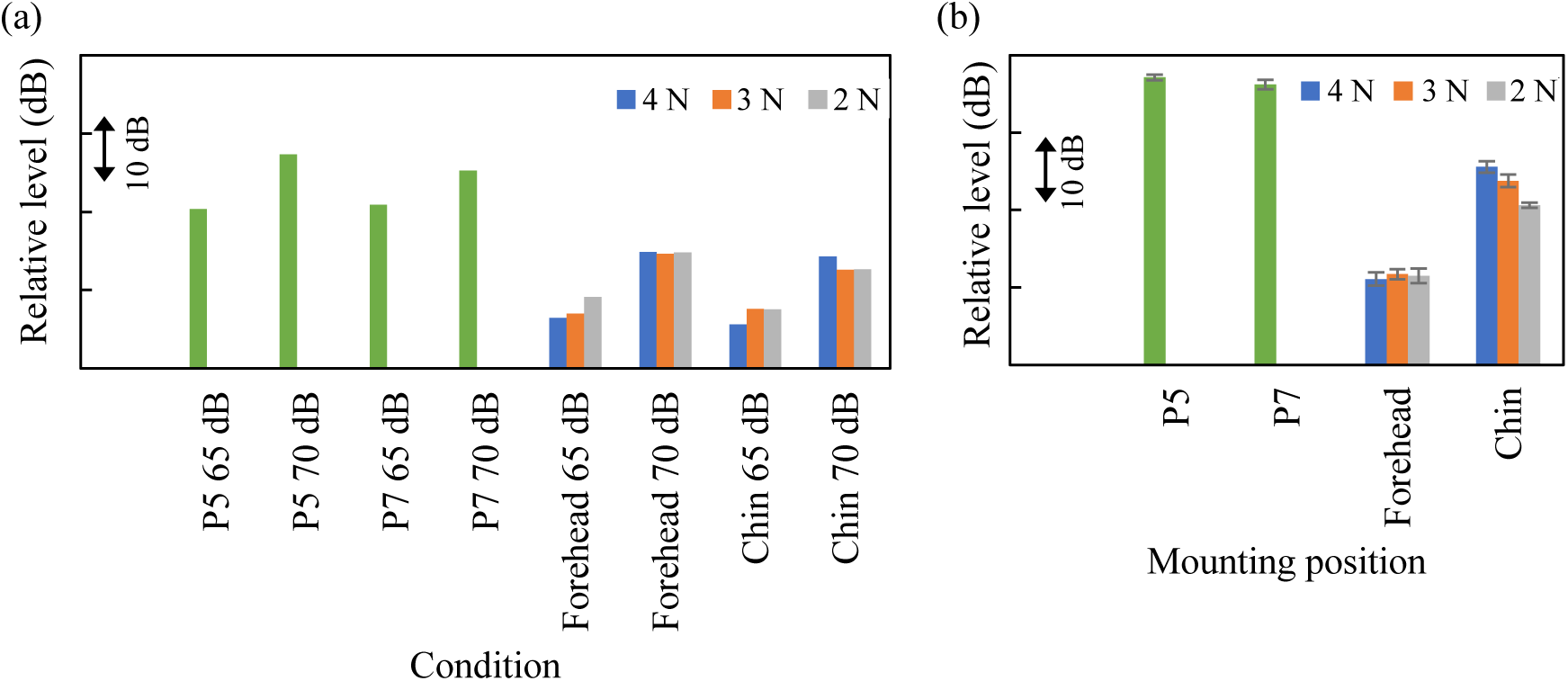
Relative sound pressure levels for each of the mounting position and reproduced background sound pressure level conditions measured in (a) background sounds and (b) speech, respectively.

### Limitations and future work

This study has some limitations. First, the present experiment examined different mounting conditions of contact microphones on the head and neck in a sitting position. However, because microphones are expected to be used during walking in a real environment, the effects of vibrations caused by walking on speech recognition accuracy should also be examined.

Second, as factors affecting recognition accuracy, this study examined speech recognition systems, gender, mounting positions, and contact force. However, there may be other important factors that were not controlled in this study, so it is necessary to explore further the factors that are important for the efficient acquisition of bone-conducted speech in future research.

Third, there may be an age bias for the subjects in the present experiment. As mentioned above, physical characteristics such as subcutaneous fat and skeletal structure may have a significant effect on recognition accuracy, so it is also important to determine how physical changes associated with aging affect recognition accuracy. Thus, experiments should be conducted on a sample with a wider age range.

## Conclusion

In this study, we investigated the effects of different contact microphone mounting conditions on speech recognition accuracy using three commonly available speech recognition systems with a focus on the operation of smart devices in public environments. A previous experiment conducted on the throat area with eight mounting locations confirmed that not only the mounting position, but also gender, affected recognition accuracy. However, in the present study, the results of the experiment on the head surface with seven mounting positions and three contact force conditions showed no effect of contact force or gender on recognition accuracy. On the other hand, the findings also suggested that the bone-conducted speech observed at the forehead and chin had smaller CERs in common with all speech recognition systems.

The results regarding the speech recognition accuracy of these two mounting positions under a simulated environment with background sounds showed that the SN ratios decreased at the forehead because the speech signal was relatively small compared with the background sounds, and that the CERs increased with increasing SPLs of the background sounds. On the other hand, the CERs did not increase with increasing SPLs of the background sounds at the chin and other positions on the throat. Therefore, it was confirmed that the bone-conducted speech obtained at the chin and other specific points around the anteroposterior positions of the mandibular angle had high noise resistance performance. Regarding the effects of the contact forces of the contact microphones on the head, the results of a subjective evaluation showed no significant difference in pain, suffering, or discomfort depending on the contact force and mounting position. Furthermore, the effects of the contact force on speech recognition accuracy were not significant. Therefore, users can choose any contact force when wearing a microphone on the head.

